# Molecular Simulations Meet Personalized Medicine. The Mechanism of Action of ClC-5 Antiporter and the Origin of Dent’s Disease

**DOI:** 10.1101/2024.11.06.622245

**Authors:** Veronica Macaluso, Carles Pérez, Robert Soliva, Yvonne Westermaier, Lucía Díaz, Miłosz Wieczór, Modesto Orozco

## Abstract

ClC-5 is a Cl−/H+ antiporter crucial for the homeostasis of the entire organism, and whose functional deficiencies cause pathologies such as Dent’s disease, a rare genetic disorder that can have lethal consequences. While the clinical aspects of the pathology are known, its molecular basis is elusive, which hampers the development of potential therapies. We present here a systematic study, where we explore the mechanism of transport of ClC-5, deciphering the choreography of structural changes required for the transport of chloride ions and protons in opposing directions. Once the mechanism is determined, we explore how the 523ΔVal deletion linked to Dent’s disease hampers the correct functioning of the transporter, despite having a very minor structural impact. Our study highlights how state-of-the-art simulation methods can provide information on the origin of rare diseases and serve as a tool in personalized medicine.

## INTRODUCTION

Dent’s disease (DD) type 1 is an ultra-rare X-linked disorder characterized by low-molecular-weight proteinuria (LMWP), hypercalciuria, nephrocalcinosis and/or nephrolithiasis, which frequently progresses to chronic kidney disease (CKD)^1^. While DD patients receive treatment to alleviate and prevent symptoms, there are no curative treatments for the disease. Most cases of DD are linked to alterations of the CLCN5 gene (Xp11.22) coding for the ClC-5 protein, a Cl^−^/H^+^ antiporter^2,3^ expressed in kidney and intestine epithelia^1^. In the human kidney, ClC-5 is found in proximal tubule cells (PTCs), co-localizing with H^+^/ATPase in early endosomes, where both regulate acidification. To a lesser extent, ClC-5 is also present in the plasma membranes (PM), where it mediates PM chloride currents and/or participates in the endocytosis of low molecular weight proteins^2,3^. Genomic studies have shown hundreds of ClC-5 variants with potential pathological profile, the most frequent being missense (35%), frameshift (31%), nonsense (16%), splicing mutations (10%), and large deletions (4%), while lower-frequency variations (1–2%) include in-frame deletions, complex mutations, Alu insertions and 5′UTR mutations^4^. Functionally, some of them are related to incorrect folding and degradation in the endoplasmic reticulum, while others lead to defective mutants unable to generate chloride currents^2,3^.

ClC-5 is part of the larger family of ClC channels and antiporters responsible for the transport of chloride, or similar anions, across the membrane. ClC channels catalyze the passive transport of the anions along their electrochemical gradients, while antiporters, such as ClC-5, use the concentration gradient of one ion to transport another one in the opposite direction with a fixed stoichiometry of 2 anions per 1 proton^1,5^. No human transporters have been experimentally characterized, but comparison of *algae* transporter, prokaryote transporters and mammal channels (PDB codes EcCLc, StCl, CmCLC, CLC-K and ClC-1) suggests that despite the different permeation mechanisms, ClC channels and antiporters show large structural similarity^6–10^. Thus, all of them are pseudo-symmetric homodimers, where each monomer comprises a trans-membrane (TM) domain, while in eukaryotes, an additional cytosolic cystathionine beta-synthase (CBS) domain is present. Each TM domain consists of 18 antiparallel α-helices, comprising an ample vestibule at either side of the membrane and a narrow pore, where two glutamic acids act as gates for anion uptake^9,11^.

While the Cl^−^ channel is mostly passive during transport, significant pH-dependent conformational changes are expected for the antiporter^12–15^. Thus, a variety of studies have highlighted the coupling of ion transport with the conformational and protonation states of the gating glutamates (Glu_ext_ and Glu_int_ pointing towards the extracellular and intracellular sides respectively)^13,16–18^, as well as with significant conformational rearrangements in the transmembrane helices affecting the the Cl^−^ pathway^13,15^. Past studies in model organisms also demonstrated that small changes away from the main translocation pathway can produce orders-of-magnitude variation in antiport kinetics^19^. However, despite all the effort focused on this system^8,13,17,20,21^, the precise sequence of events responsible for ions passage is still unclear, hampering the possibility to define the molecular origin of the pathological nature of certain genomic alterations, like those responsible for DD.

In this contribution, we describe the mechanism of ion transport of ClC-5 and how it is altered by the ClC-5 523ΔVal in-frame deletion, a poorly characterized rare genetic alteration associated with DD^2,3,22,23^, for which a knock-in mouse model is available^24^. By combining a large variety of *state-of-the-art* simulation techniques on the wild type (WT) and 523ΔVal variant, we obtain a detailed model of the 2:1 antiport mechanism, deciphering a series of coordinated movements coupled to the inverse flux of Cl^−^ and H^+^. By comparing the transport mechanism of the wild type (WT) and the 523ΔVal variant, we define the molecular basis of the 523ΔVal pathogenicity, and highlight some structural differences which might be used to design drugs that recover the normal functionality of the antiporter.

## METHODS

### Structure preparation and Molecular Dynamics (MD) simulations setup

As no structure of human mammal antiporters is available, we tried different homology modelling strategies to obtain structures for WT and 523ΔVal forms of the ClC-5 transporter, eventually settling on AlphaFold-Multimer predictions^25–27^ as implemented in COSMIC2^28^. Predicted structures show high-confidence scores and the expected fold for an antiporter (very close to the algae structure; see Figure 1). Each of the ClC-5 monomer comprises the trans-membrane (TM) and the cytosolic cystathionine beta-synthase (CBS) domains. Missing hydrogens were added using the H^++^ webserver^29^, according to their pKa values. By default, the glutamate residue 211 (Glu_ext_) was protonated in both monomers, while glutamate residue 268 (Glu_int_) was considered deprotonated. As discussed later, this would correspond to the starting configuration for chloride transport from the cytoplasm to the intracellular space.

**Figure 1:**
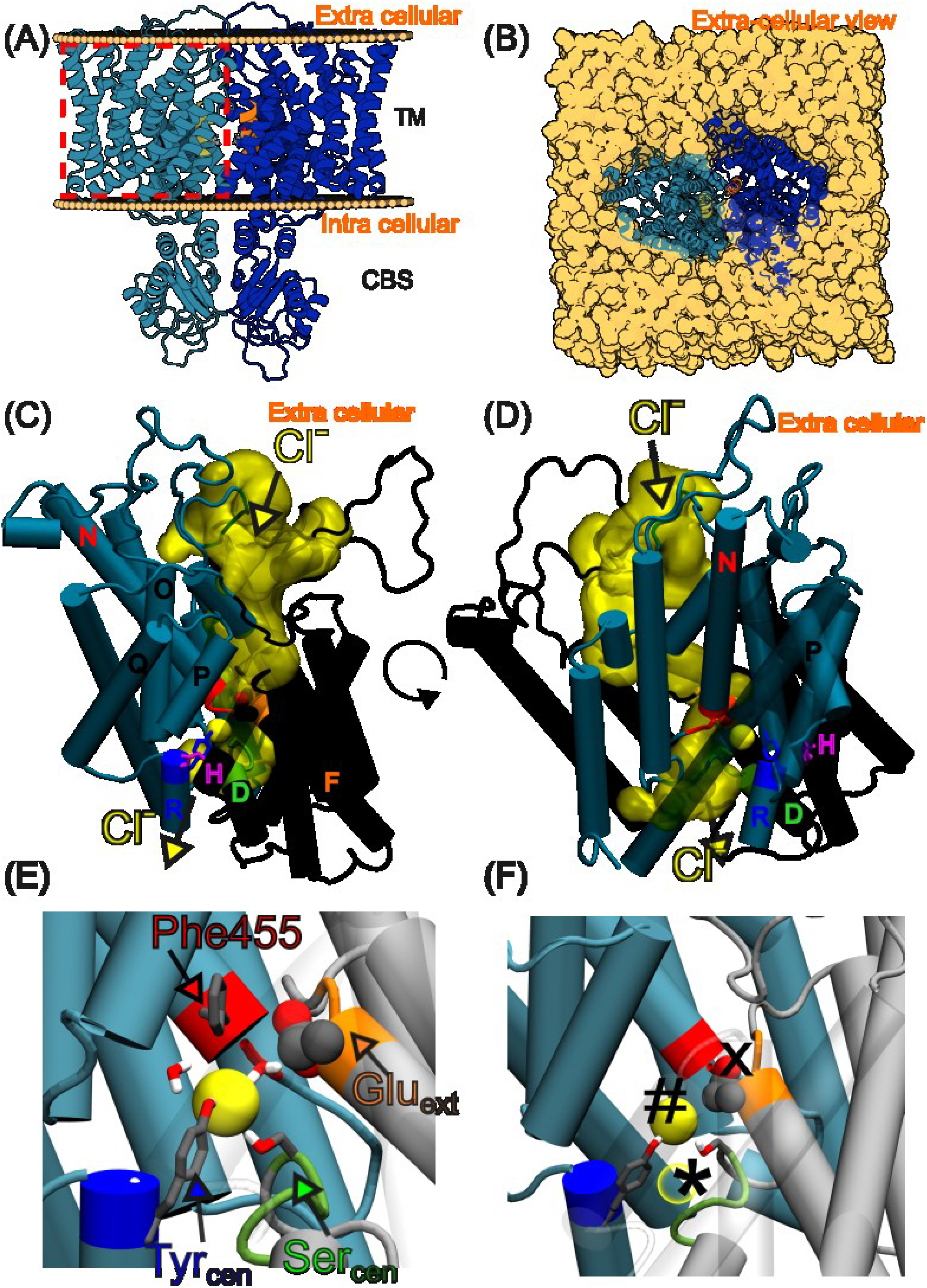
Wild-type ClC-5 homodimer (A, B), details of the TM domain (C, D), and binding sites representation (E, F) in two side views. Panel A and B show side and top (extra-cellular) view of wild-type ClC-5, respectively. Chains A and B are in blue and dark blue, with the α-helix P (where Val523 is placed) is yellow (chain A) or orange (chain B). Panels C, D, E and F show details of one of the wild-type monomers, with α-helices A to I in gray, and α-helices J to R in cyan. Electropositive N-termini of α-helices D (green), F (orange) and N (red) and key residues for the ClC-5 antiporter mechanism are highlighted (Tyr_cen_, Ser_cen_ and Glu_int_ licorized blue and green and magenta respectively and Glu_ext_ in CPK; global pictures are shown in panels C and D and more detailed representation are shown in panels E and F). Yellow arrows indicate Cl^−^ ions pathway. Panels E and F show a molecular representation of chloride binding sites: **S**_**ext**_ (labeled with “X”) is occupied by Glu_ext_ side chain; **S**_**cen**_ (labeled with “#”) is occupied by a chloride anion (yellow solid sphere); and **S**_**int**_ (labelled with “*”) which can be occupied by another chloride (yellow empty sphere). The detailed placement of the Val523 in the structure (helix P) is shown in Suppl. Figure S2.

The protein structures yielded by AlphaFold were embedded in a phosphatidylcholine (POPC) bilayer and immersed in a rectangular water box (13 × 13 × 15 nm and 13 × 13 × 14 nm dimension for WT and 523ΔVal, respectively) using CHARMM-GUI Membrane Builder^30^. To favor transport, chloride ions (Cl^−^) were added to reach 1 Molar concentration, while sodium ions (Na^+^) were added as needed to charge-neutralize the system. CHARMM36 *force-field*^*31*^ was used to model protein, anions, and lipids, while water was modelled using TIP3P^32^. Systems (containing around 240,000 atoms) were minimized, thermalized and equilibrated before performing production runs (see Suppl. Methods for details). Small local distortions found in the AlphaFold model were quickly relaxed during equilibration, yielding very stable structures (see Suppl. Figure S1). Equilibrium trajectories of 3 replicas were performed for 1 μs per system (WT or 523ΔVal) using GROMACS2020 software (see Suppl. Methods for additional details)^33^.

### Tunnels identification

CAVER^34^ was used to identify tunnels in the equilibrated structures of WT and 523ΔVal. Representative snapshots for the tunnels analysis were obtained by a *k-means* clustering analysis using the RMSD of residues in proximity of Glu_ext,_ Ser_cen_ and Tyr_cen_ as a descriptor (see Figure 1). To define “productive cavities”, we imposed a condition that tunnels should connect from the bulk (outer surface of the protein at the intracellular or extracellular space) to the external-gate region.

### Multiple-walker metadynamics and pKa calculations

Multiple-walker well-tempered metadynamics (MWWTMetaD)^35,36^ was used to explore the correlated free-energy surface of two chloride ions confined in the central channel. Eight simulations were run for each monomer of the WT and 523ΔVal variant dimers for two different protonation states of the gating glutamates (Glu_ext_ protonated and Glu_int_ deprotonated, and Glu_ext_ deprotonated and Glu_int_ protonated). Starting configurations were selected from the ensembles obtained by unbiased molecular dynamics, where the density of Cl^−^ around the entrance of the anion tunnel was high. This process led to 64 independent walkers (8 walkers x 2 monomers x 2 protonation states x 2 protein forms), symmetrized and combined to obtain 4 free energy landscapes of the (permutation-symmetric) chloride pairs, named **WP** (wild type with the Glu_ext_ protonated and Glu_int_ deprotonated state), **WD** (wild type with Glu_ext_ deprotonated and Glu_int_ protonated), **MP** (the 523ΔVal mutant form with the Glu_ext_ protonated and Glu_int_ deprotonated state), and **MD** (the 523ΔVal mutant form with Glu_ext_ deprotonated and Glu_int_ protonated). In all cases, we used two collective variables defining the Z-displacement (i.e. along membrane normal) of the two chlorides from the center of mass of the central cavity (see Suppl. Methods for details). To accelerate convergence, after ca. 1 μs per walker, we averaged and symmetrized hills for both monomers, and restarted the simulations with a 4-fold larger initial hill height for another >500 ns per walker. Details of the associated simulations were identical to those used to obtain the unbiased trajectories.

### Effective pKa calculations

We used classical molecular interaction potential (CMIP)^37^ (with a non-linear Poisson-Boltzmann’s electrostatic term; see Suppl. Methods) to compute configuration-dependent pH for the two gating glutamic acids. To this end, metadynamics trajectories were binned according to the Z position of both chloride ions, and the calculation was performed on up to 10 selected frames with highest statistical weights in each bin (a total of 12906 structures) according to the metadynamics reweighting scheme^38^; the pKas corresponding to individual frames were then averaged. In all cases, CHARMM36 charges and van der Waals parameters were used in CMIP combined with standard dielectric constant for protein and solvent and physiological ionic strength for the Poisson-Boltzmann part of the calculation.

### Determination of druggable cavities

Preliminary druggability studies were performed using Fpocket^39^ on 10 selected structure representatives of highly populated clusters from the unbiased simulation of the 523ΔVal variant. A more exhaustive analysis was done using MDpocket^40^, a method that allows us to account for cavity plasticity using unbiased trajectories of both WT and 523ΔVal variants (see Suppl. Methods for details).

## RESULTS AND DISCUSSION

### AlphaFold model and solution structure

Consistently with known structures of other ClC proteins, the structural model of ClC-5 generated by AlphaFold and refined by molecular dynamics simulations shows a dimeric protein (Figure 1A and B), where each TM domain consist of 18 α-helices. An ample vestibule (Figure 1C and D) is present at both sides of the membrane, and a narrow pore, whose center is defined by the N-terminal regions of α-helices D, F and N, and an R-helix Tyrosine (Tyr_cen_), which generate a central cavity (**S**_**cen**_). This cavity acts as an anion trap, and is separated from the cytoplasmic side by the side chains of D-helix Ser_cen_ and R-helix Tyr_cen_. Moving closer to the extracellular side, another cavity, hereafter named **S**_**ext**_, is located between the α-helices N and F, and is able to accommodate either the negatively charged side chain of an F-helix Glutamate residue (Glu_ext_) or a chloride anion. Finally, a third cavity **S**_**int**_ is located below **S**_**cen**_, flanked by the N-terminus of α-helix D on the one side, and the intracellular aqueous vestibule on the other. This cavity is expected to be important for the recruiting of ions from the intracellular environment, or in the chloride exit pathway in the opposite direction^20^ (see Figure 1 for description of the crucial regions of the anion tunnel). These three cavities are in very close proximity, defining a narrow gating site that controls the flux of ions in the channel.

According to structures refined through molecular dynamics, the 523ΔVal deletion leads to minor structural changes: the only noticeable change is a certain compaction between the two subunits (see Figure S1). The deletion is well accommodated, maintaining the secondary structure on helix P until Arg516, where a short turn recovers the phase of the helix without major changes in the contact residues (see Figure S2). CAVER analysis (see Methods) shows the existence of four tunnels connecting the extra-cellular or intra-cellular vestibule to Glu_ext_, which can be assigned to the preferred H^+^ and Cl^−^ paths (Figure 2) and which are all present in both WT and 523ΔVal variants.

**Figure 2:**
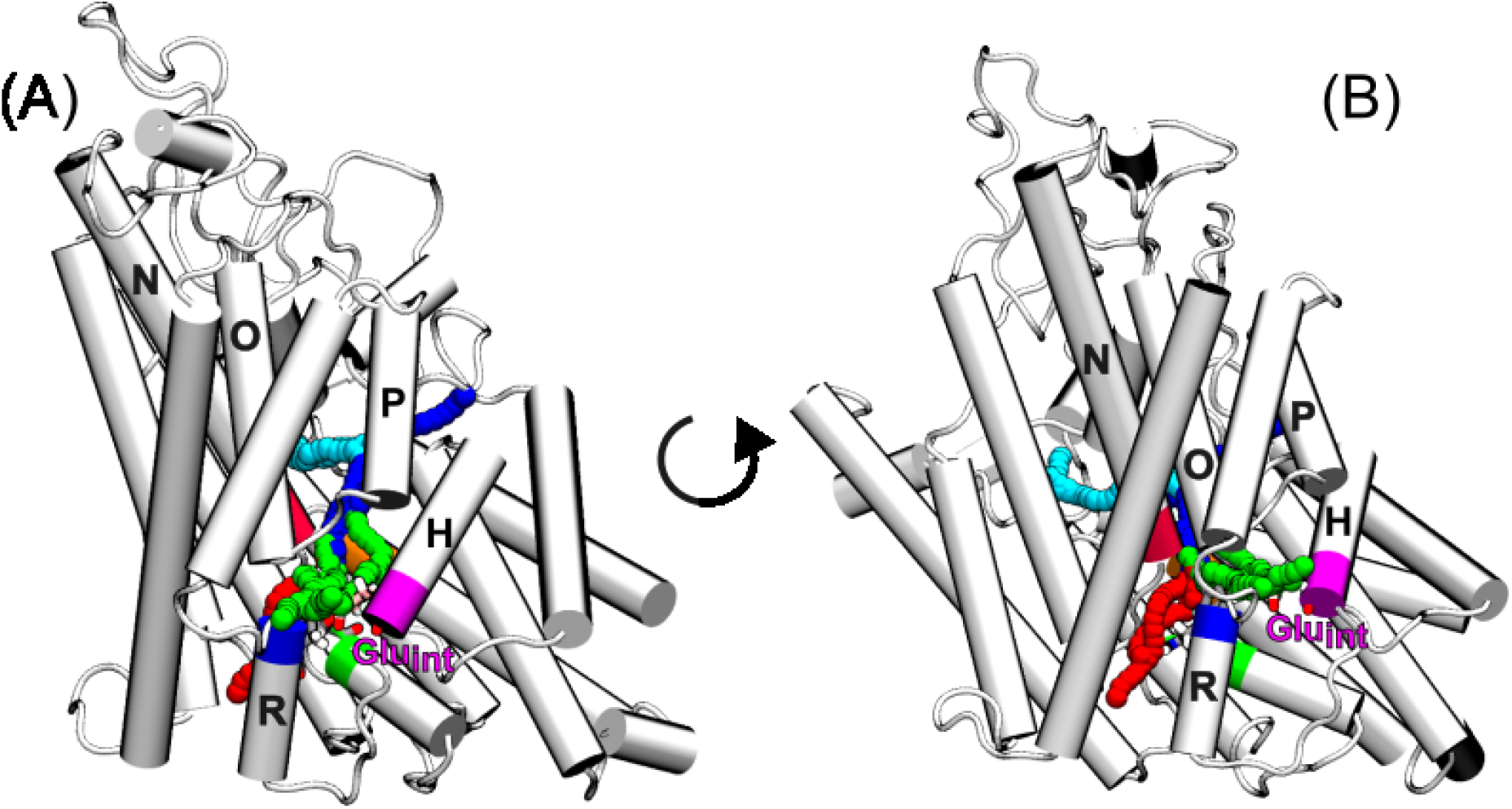
Visualization of the four tunnel clusters obtained for Wild Type and 523ΔVal systems projected on a representative molecular structure of Wild Type TM chain. Figure A and B offer a different-rotation lateral view of the TM chain. α-helix H, on which Glu_int_ is located, is colored in magenta. The cyan and blue tunnels start from the extra-cellular vestibule and resemble the extra-cellular Cl^−^ and H^+^ common path described in the literature for other ClC channels. The red tunnel cluster goes from the intra-cellular vestibule (figure bottom) and passes in proximity of Tyr_cen_ and Ser_cen,_ representing the intra-cellular chloride path; the green one starts from the intra-cellular vestibule near Glu_int_ and represents the intracellular H^+^ path.

### Thermodynamics of the mechanism of Cl^−^ transport

#### The chloride migration

Free energy surfaces obtained with MWWTMetaD (see Methods) allowed us to determine the pathway of Cl^−^ migration through the transporter (see Figure 3A). As shown in Figure 3A-WP, the first chloride ion can very easily reach a stable binding region defined by three closely placed free energy minima M1_w_ (**S**_**ext**_), M2_w_ and M3_w_ (**S**_**cen**_), the last of them acting as a kinetic gate. When the first Cl^−^ waits at **S**_**cen**_, the second ion can arrive at the extracellular vestibule, but is unable to enter further inside the channel due the presence of the first Cl^−^, which in turn cannot progress to the intracellular side due to a high free-energy barrier (Figure 3A-WP). Escape from this stable situation requires a shift in the protonation state of Glu_ext_. According to the **WT** pKa plot (see Figure 3B), this is indeed what happens: when the bottom chloride reaches S_cen_, and a thermal fluctuation moves the top chloride deeper into S_ext_, the pKa of Glu_ext_ shifts, allowing the residue to release the proton to the extracellular vestibule. A subsequent change in Glu_int_ (Figure 3B) leads to a protonation situation Glu_ext_(deprotonated)-Glu_int_(protonated), which facilitates the migration of the two Cl^−^ as shown in the **WD** free energy surface (Figure 3A-WD). During deprotonation, the top Cl^−^ ion can enter deeper into the pore, so that the M4’_w_ minimum is reached. In the free energy profiles, this corresponds to a diagonal displacement from M3_w_ (**WP**) to M4’_w_ (**WD**), from which both ions can eventually exit the pore through simple diffusion, with large free energy barriers preventing the return of the ions towards the entrance.

**Figure 3:**
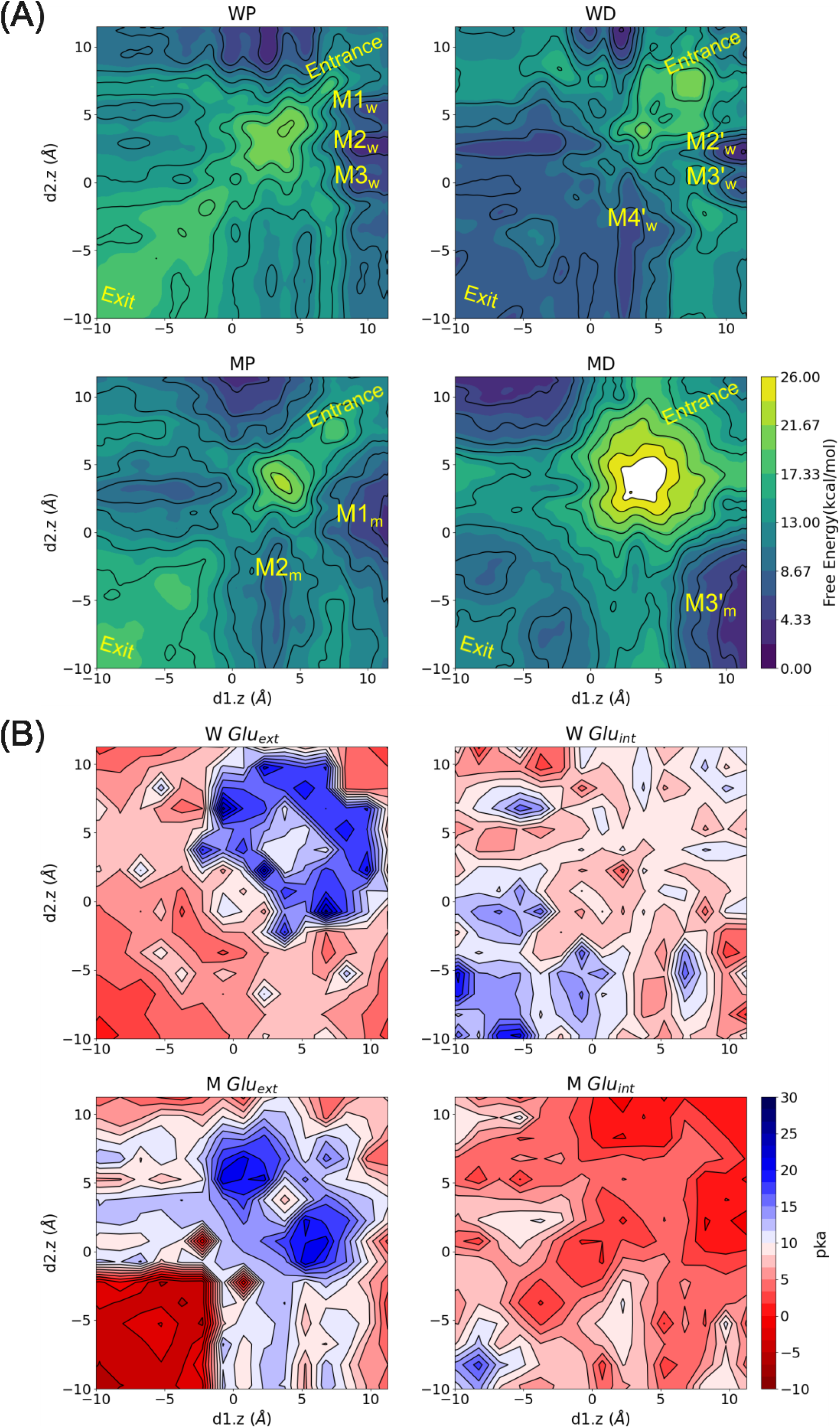
(**A**) Free energy surfaces (FES) obtained from multiple-walker metadynamics simulations of WT (WP and WD) and 523ΔVal (MP and MD) systems (where W means WT; M the 523ΔVal variant; P means that Glu_ext_ is protonated and Glu_int_ is deprotonated, and D stands for the reverse). Note that according to the defined collective variable (see Methods) the (10/10) region corresponds the two Cl-at the extracellular side and the (−10/-10) to the two Cl-at the intracellular side. Movement along either the X or Y axis represents the displacement of one Cl-from the extra (positive values) to the intra (negative values) sides; movements along the diagonal means coordinate movement of the two ions. (**B**) pKa of external Glu_ext_ and Glu_int_ for WT (W, top panels) and 523ΔVal (M, bottom panels) systems as calculated using the CMIP method.

A complementary molecular-level explanation of the chlorides’ migration is shown in Figure 4. Initially, extracellular chloride ions are attracted by backbone atoms of loops L1-L2 and E-F, Ser380 and Thr381 of helix L2, and repelled by hydrophobic residues on helix E. After passing the positively charged Lys210 acting as an ion filter (just one residue before Glu_ext_), the bottom Cl^−^ reaches the **S**_**ext**_ cavity, a site enclosed between helices N and F, and protected from extra- and intracellular sides by loops E-F and M-N. At the same time, the other ion can reach the outer side of the channel (**S**_**out**_). This defines the M1_w_ state, a minimum in the free energy surface in Figure 3A. At the second minimum M2_w_, the E-F loop assumes a new conformation, widening the distance between helices F and N. This allows the descent of the Cl^−^ dwelling at S_ext_ to reach the S_cen_ cavity in the neighboring state M3_w_, while the other Cl^−^ remains at S_out_, enlarging the separation between the two chlorides. The **S**_**cen**_ binding site is defined by conserved Ser168 (Ser_cen_), Tyr558 (Tyr_cen_), Leu454, Ile170, Phe446 and Leu454 and holds the ion very tightly, hindering the progress of the ion towards the exit. This stable binding site can only be disrupted by the change in the protonation state of Glu_ext_ predicted by pKa maps in Figure 3, as its side chain reaches the water molecules above the top chloride. With the Glu_ext_ deprotonated, the second chloride is forced into the channel reaching state M4’_w_, followed by a spontaneous diffusion of the two ions through a low stability **S**_**int**_ binding site (Figure 4). In parallel, the now favorable protonation of Glu_int_ prepares the cycle for regeneration.

**Figure 4:**
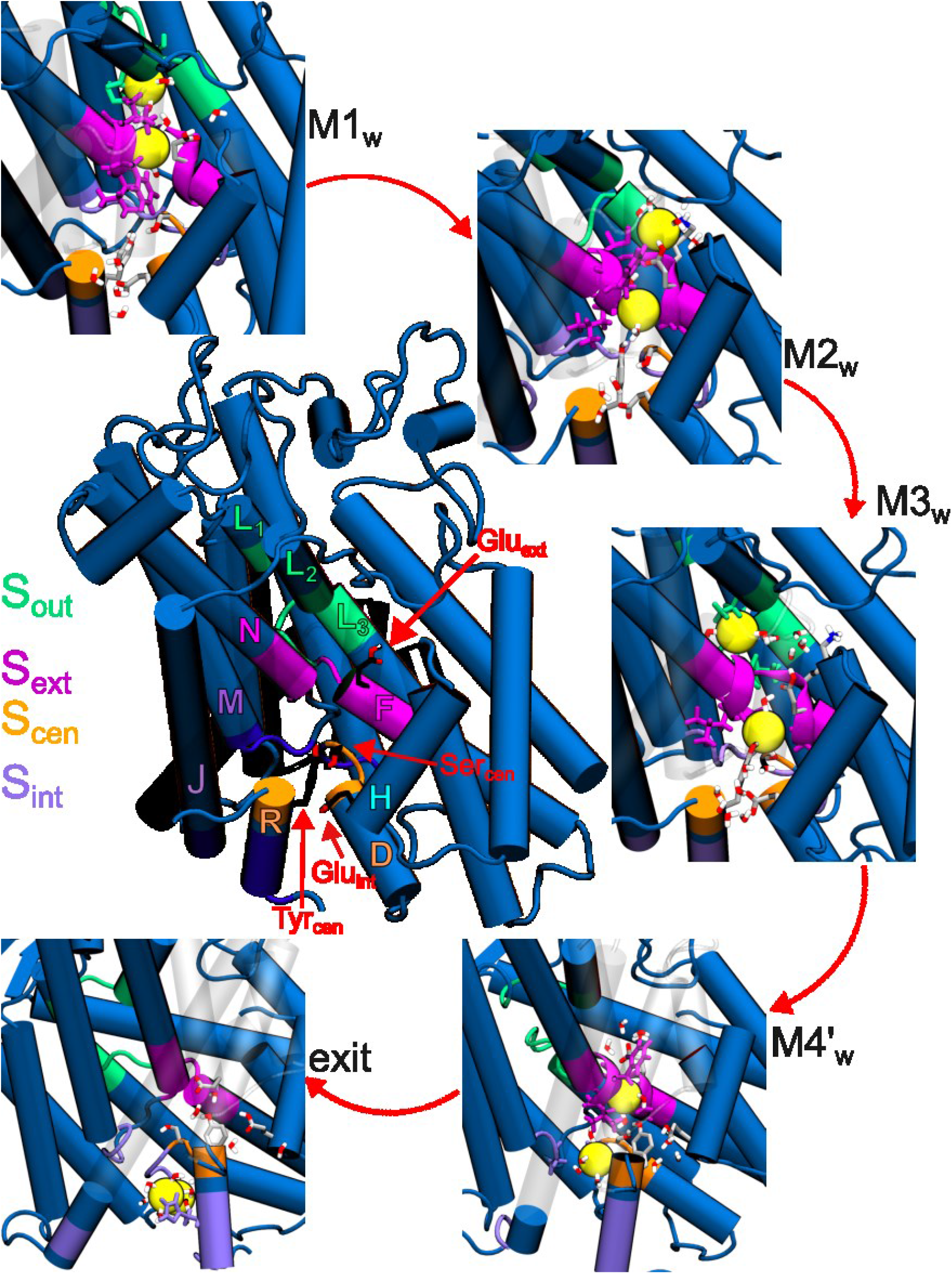
Molecular representation of mw-MD FES minima and anion binding sites of wild-type ClC-5. In the central panel we represent the location of four identified chloride biding sites in a TM chain. The other panels show a molecular representation of WP (M1_w,_ M2_w_ and M3_w_) and WD (M4_w_ and exit) minima.

#### The proton migration

The mechanism outlined above explains the transport of two Cl^−^ from external to intracellular spaces at the expense of a switch in the protonation state of Glu_ext_ and Glu_int_, yielding a net proton transfer from intracellular space to the exterior. Note that when the Cl^−^ transport is completed the protonation state is: Glu_ext_-protonated and Glu_int_-protonated state, a configuration that would allow for reverse transport from the intracellular space to the cytoplasm (see Figure 3, WD and WP). This would in turn lead to a futile cycle without a net flux of Cl^−^. However, if the concentration difference of Cl^−^ and/or H^+^ between the two sides of the membrane prevents it, the net flux of ions will continue in the same direction. For the transporter to be regenerated, the proton that was originally transferred from the intracellular space to Glu_int_ must move to Glu_ext_. An analysis of the trajectories show that this regeneration can happen through a Grotthus’ mechanism, taking advantage of a water spine connecting both glutamic acids when anions are reaching the intracellular vestibule and in equilibrium simulations, i.e., when the respective pKa values favor the exchange (see Figure 5).

**Figure 5:**
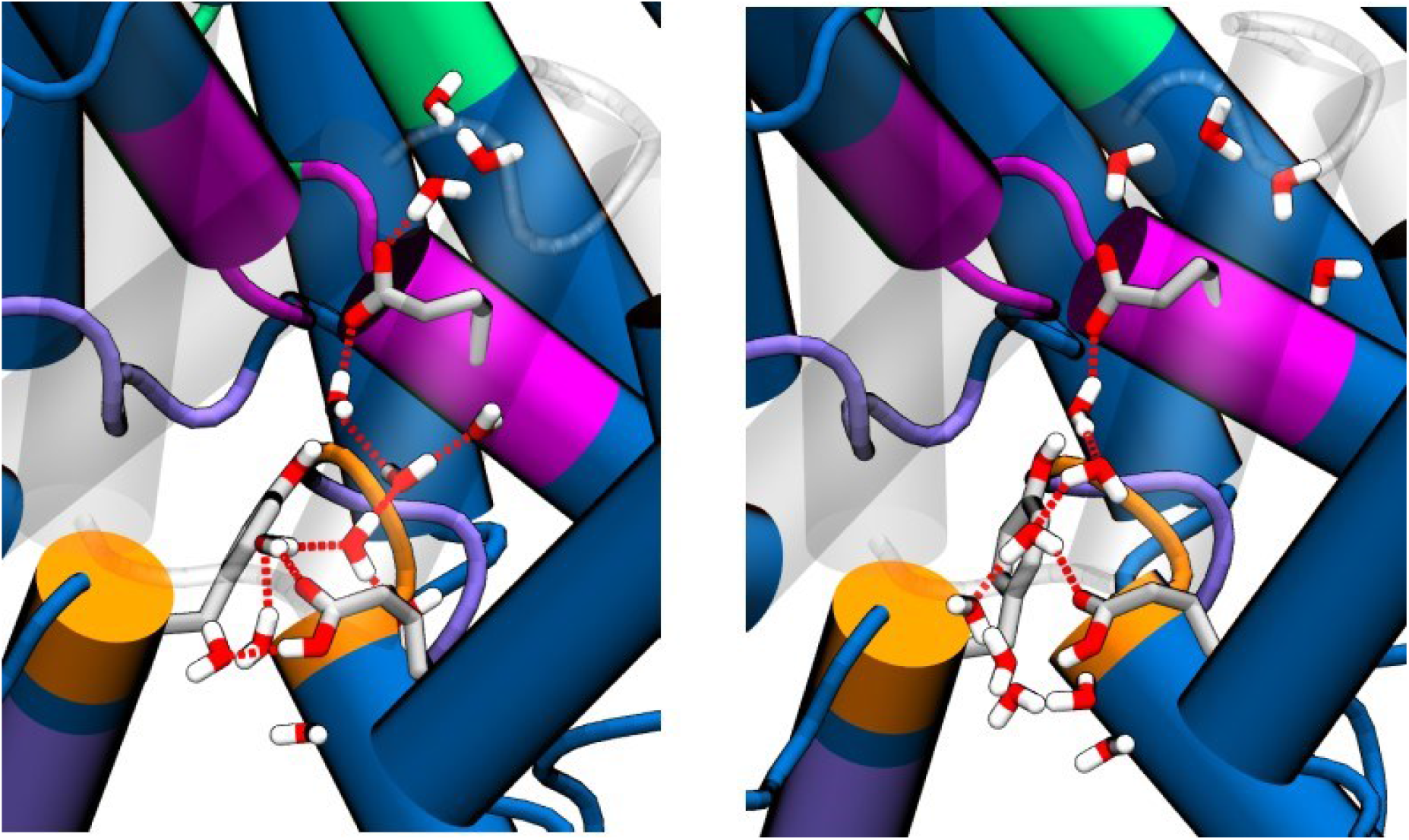
Two examples of a water wire connecting Glu_ext_-deprotonated and Glu_int_-protonated which would justify regeneration of the enzyme by recovering the Glu_ext_-protonated and Glu_int_-deprotonated state.

### The impact of the deletion of valine 523

The deletion of Val523 leads to minor changes in the structure: a moderate increase in inter-subunit compaction (Figure S1A) and a slight displacement of helix P (Figure 2, S1C, and S2). The impact of such structural changes on the antiporter mechanism is however dramatic (Figure 3), consistently with the clinical impact of the mutation. As shown by the MWWTMetaD simulations, the simultaneous transfer of two Cl^−^ as in the WT protein is forbidden (diagonal movement in MP plot in Figure 3A), but a single Cl^−^ can enter inside the channel so that a state similar to M3_w_ is reached, provided that the external glutamate is in the protonated state. The differences arise when the external glutamate releases a proton to the extracellular vestibule and the internal one takes one from the internal vestibule (i.e. globally Glu_ext_H--> Glu_ext_^−^ and Glu_int_^−^ →Glu_int_H). In the case of the WT, this change in protonation states of the glutamates leads to a barrierless path for the simultaneous transfer of two Cl^−^ from M3’ to M4’ and the exit (see the favorable free energy values in the left bottom part of WD plot in Figure 3A). In turn, the change in protonation states of the two glutamates in 523ΔVal variant leads to a free energy surface where the coordinated transport of two chlorides is forbidden by large free energy barriers, favoring the transfer of a single Cl^−^ ion through the M3’m state and its subsequent diffusion to the intracellular vestibule. This would suggest a one-to-one stoichiometry and an inferior ability to maintain homeostasis. Furthermore, analysis of the pKa plots (Figure 3B) for the different structures collected in the metadynamics simulations show interesting differences between the WT and deletion variants. As described above, in the WT protein the entrance of Cl^−^ into the channel is incompatible with Glu_ext_ in the deprotonated form, and the same holds true for the 523ΔVal variant (see WT Glu_ext_ and 523ΔVal Glu_ext_). When the two Cl^−^ are reaching the exit, Glu_ext_ becomes very acidic (and accordingly deprotonates) in both WT and deletion variant, even more in the second case (see Suppl. Figure S3). Major differences appear however for Glu_int_ between the WT, where it can easily protonate when one Cl^−^ is inside the channel (see blue surfaces in the bottom-left part of the W Glu_int_ plot in Figure 3B), and the 523ΔVal variant, where very low pKa values hinder protonation when one or even two chlorides are forced to be in the channel (see M Glu_int_ plot in Figure 3B). In other terms, the change in the protonation state of Glu_int_ that is required for the two chlorides to reach the exit cavity is much more difficult in the 523ΔVal variant, reinforcing the problems for dual chloride transport.

The structural analysis helps us rationalize the changes induced by an apparently mild deletion that seems to be locally well tolerated (see Suppl. Figures S1-S2). Indeed, the movement of helix P resulting from the deletion leads to (1) altering the local environment of the Glu_int_ (located at the neighboring helix H), which becomes close to Lys174 on helix D, (2) dehydrating the acidic group, and (3) stabilizing the anionic state (see Suppl. Figure S4). Analysis of the structures around the crucial position (d1.z=3,d2.z=−5) in Figure 3 shows the impact of the deletion on the tilting of the P helix. This in turn has two major effects: the loss of a close electrostatic interaction stabilizing one of the chloride ions at S_cen_ in the WT but not the deletion variant (gray arrow in Figure 6), and the generation of a steric clash for the advance of that chloride in the deletion variant (orange arrow in Figure 6). Thus, the M4 minimum disappears in the 523ΔVal variant, excluding the possibility for the two ions to reach simultaneously the intracellular vestibule.

**Figure 6:**
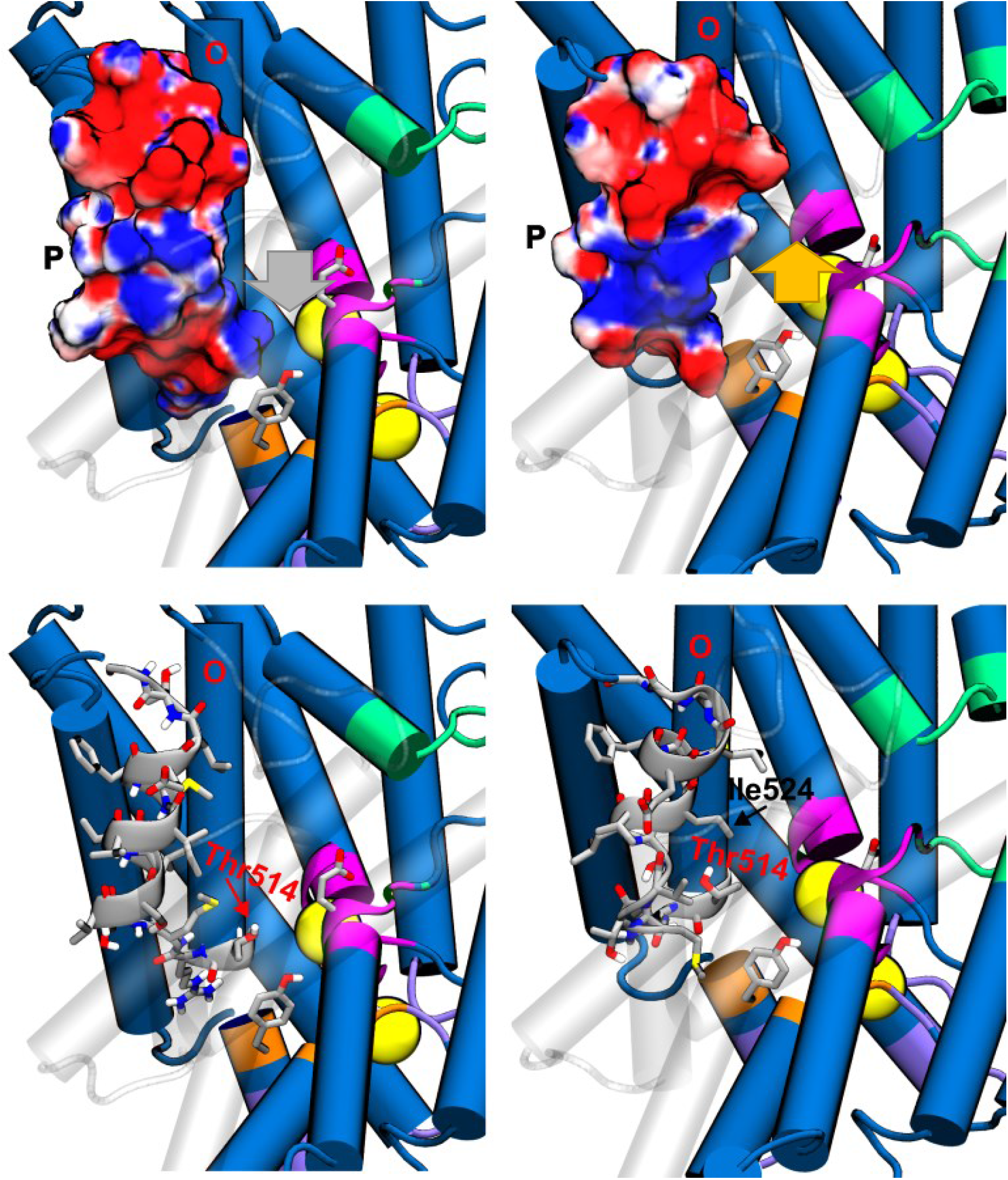
Structural details of the channel in the M4’_w_ region (see Figure 2) which correspond to a free energy minimum for the WT (left column), but not for the deletion variant (right column). Top row shows electrostatic surfaces, while the bottom row shows side details. Chlorides are shown as yellow spheres.

Finally, we explored potential changes in drug-binding properties emerging from the deletion of valine 523. To this end, we ran Fpocket calculations (see Methods) to detect drug-like cavities in WT and 523ΔVal variant. Interestingly, a sizeable cavity with properties that would allow binding of a drug was detected in the vicinities of helix P (see Figure 7). Fpocket calculations suggest that such a cavity is sensitive to the deletion of the valine 523 (Figure 7A-C), a result that is confirmed by dynamic calculations using MDpocket (see Methods and Figure 7D-F), which highlights the presence of variant-specific sub-pockets. Such features could be exploited in the future to develop pharmacological chaperones interacting with the pathological variant to recover the correct alignment of helices P and H, or to more directly modulate the pKa of Glu_int_.

**Figure 7:**
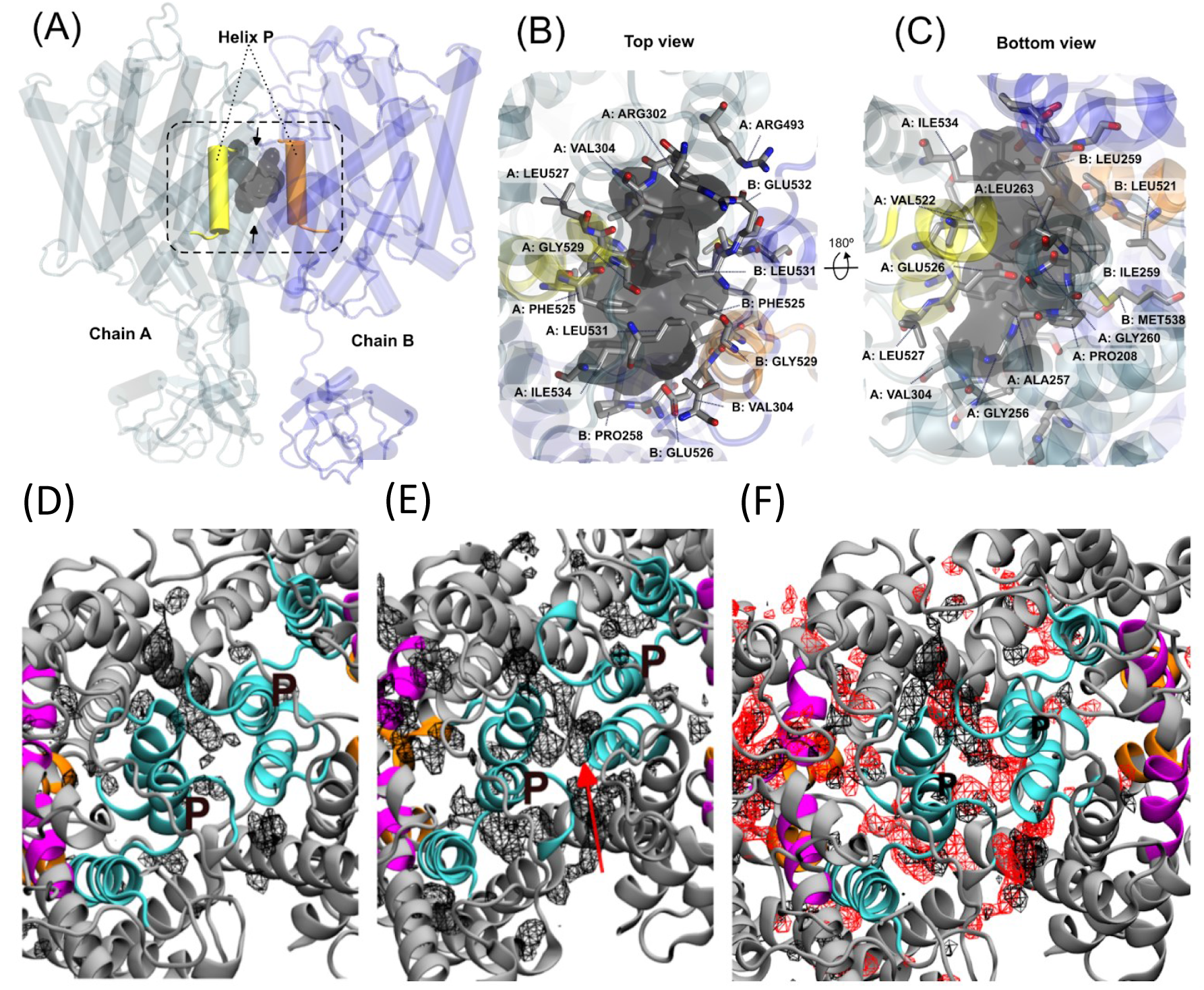
First row: druggable pocket of the representative cluster of the MD for the ValΔ523 mutant variant of ClC-5 homodimer. (A, B, C). Calculations were perfomed using Fpocket on 10 representative structures from the deletion variant (Druggablity score: 0.990. The homodimer is represented in two chains: blue for A and dark blue for B. The helix P of chain A and chain B are colored in yellow and orange respectively. The druggable site candidate is shown with a dark grey surface. (A) Global point of view of the druggable site, located in between the two helixes P. The arrows represent the different points of view described in (B) and (C). (B) Zoom in into the druggable site, view from the top, and (C) from the bottom. Second row: MDpocket density maps for WT and 523ΔVal variants. Relevant channel helixes are in color cyan (helix P labeled), pink, and orange in (D) and (E) the density maps are shown in black wires. (D) Density map of the WT protein, using a threshold of 5.7, while in (E) the 523ΔVal variant the threshold is 3.3. The red arrow shows a subpocket only present in the 523ΔVal variant. (F) Superimposition of (A) and (D), with WT in black and 523ΔVal in red.

## CONCLUSIONS

The transfer of Cl^−^ in antiporters requires a shift in the protonation state of two conserved glutamates located at the two ends of a narrow tunnel controlled by up to four cavities with high residence times for the Cl^−^. The central site acts as a trap for one Cl^−^ which blocks the entrance of the second one. Escape from this site requires a coordinated change in the protonation state of the external glutamate that interchanges protons with the extracellular vestibule, triggered by the presence of the second chloride. The shift in protonation states leads to a downhill free energy landscape where two chlorides can migrate almost simultaneously, as well as allows to protonate the internal glutamate. This in turn brings the transporter to a final state that can be regenerated by a Grotthus mechanism involving a short chain of water molecules, with a net flux of one proton from intracellular to extracellular space. Interestingly, the entire cycle can operate without major conformational changes, relying instead on local rotameric changes of key residues.

According to our results, the mechanistic disruption caused by the 523ΔVal variant does not rely on global structural rearrangements and is unlikely to produce major changes in stability, distribution or dimerization properties, but rather induces small local changes in the vicinities of the internal Glutamate and the central site S_cen_. These apparently minor changes (1) modify the effective pKa of Glutamates, making the proton migration very difficult, and (2) change the preferred chloride migration path, making the simultaneous transfer of two chlorides impossible, and hampering a correct homeostasis in the kidney.

## ACKNOWLEDGMENTS

This work was funded by the ASDENT Foundation, the Spanish “Ministerio de Ciencia e Innovación” (PID2021-122478NB-I00), the Center of Excellence for HPC H2020 European Commision. “BioExcel-2. Centre of Excellence for Computational Biomolecular Research” [European Union: 101093290 and Ministerio de Ciencia e Innovación: PCI2022-134976-2]. This project is co-funded by the European Regional Development Fund under the framework of the ERFD Operative Programme for Catalunya, the Catalan Government AGAUR (SGR2017-134). The authors thank the Barcelona Supercomputing Center for a HPC resources. We also gratefully acknowledge Poland’s high-performance Infrastructure PLGrid ACK Cyfronet AGH for providing computer facilities and support within computational grant plgcgconf.

