## Supplementary Material for "Molecular Simulations Meet Personalized Medicine. The Mechanism of Action of ClC-5 Antiporter and the Origin of Dent’s Disease"

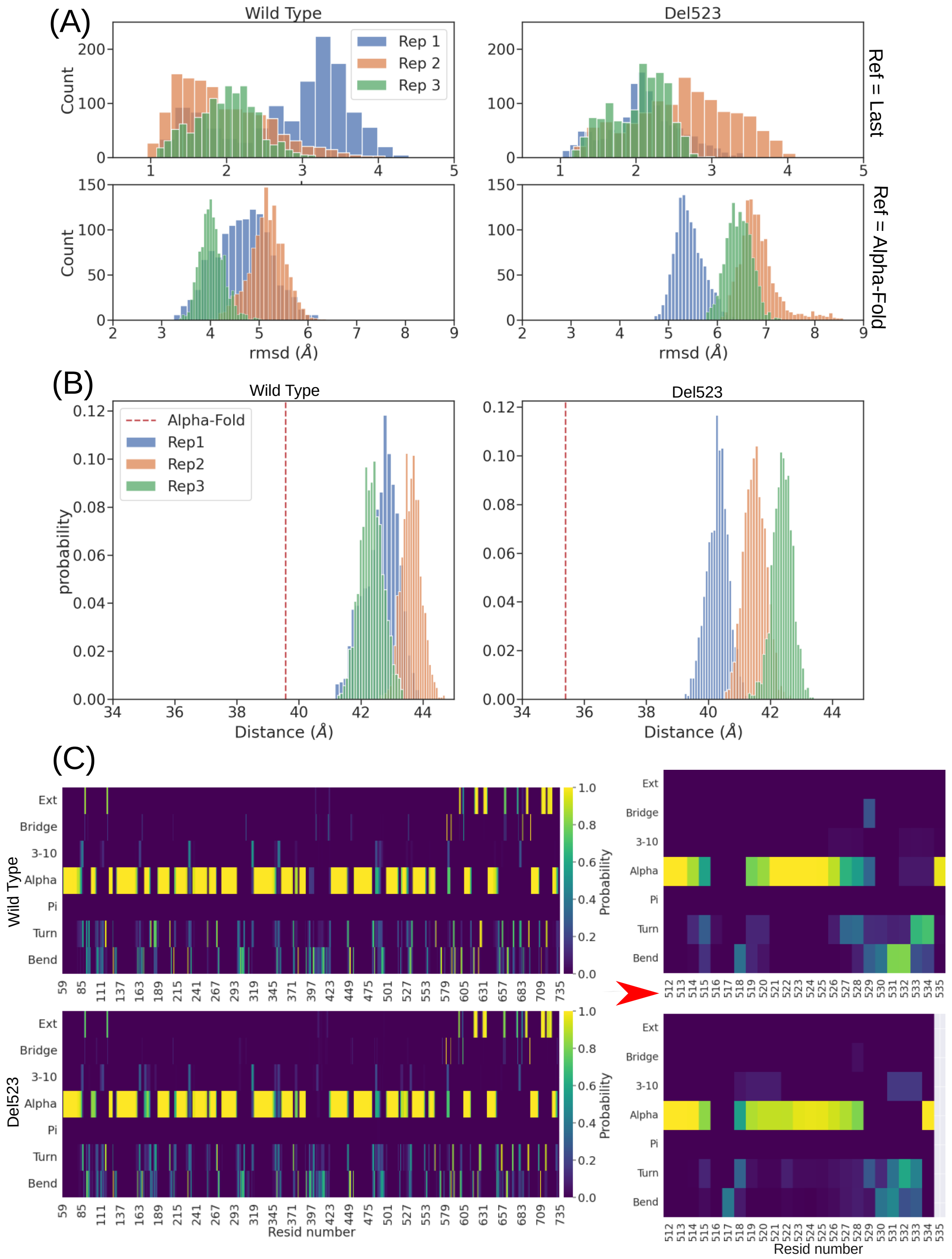


*Figure S1: (A) RMSD, (B) Center-of-mass distance and (C) secondary structure analysis carried out over WT and 523ΔVal plain MDs. (A) Backbone-atoms RMSD histograms calculated using the last 500 ns of each plain MD replicas (Rep1, Rep2 and Rep3). In the two top panels the reference is the (WT or 523ΔVal) average structure calculated over the last 100 ns of the corresponding MD replica. In the bottom panels the reference is the corresponding AlphaFold model structure. (B) Histograms of the distance between the center-of-mass (COM) of the two monomers trans-membrane unit. The COM distance for the corresponding Alpha-Fold model is also reported. (C) Secondary-structure heatmap calculated over each MD replica monomer. In the right panel a zoom on the α-helix P is shown.*

(A)

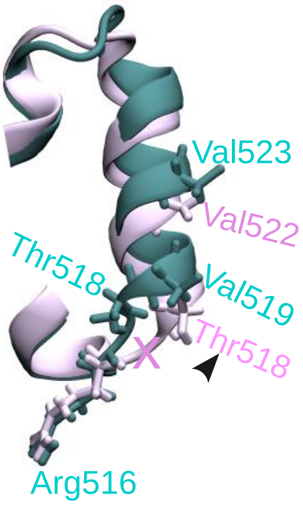


(B)

| | | | | | | | | | | | |
|---|---|---|---|---|---|---|---|---|---|---|---|
| $AA_n$ | Arg | Met | Thr | Val | Ser | Leu | **Val** | **Val** | Ile | Met | $AA_{n+1}$ |
| n | 516 | 517 | 518 | 519 | 520 | 521 | 522 | 523 | 524 | 525 | n |
| n | 516 | 517 | | 518 | 519 | 520 | 521 | 522 | 523 | 524 | n-1 |
| $AA_n$ | Arg | Met | | Thr | Val | Ser | Leu | **Val** | Ile | Met | $AA_{n+1}$ |

*Figure S2: (A) helix-P superimposition of WT and* 523ΔVal *average MD structures; (B) Alignment of WT and* 523ΔVal *residues, as observed in plain MD average structure. WT is colored in cyan and* 523ΔVal in light pink.
*Up to Arg515, WT and* 523ΔVal *perfectly align in both primary and secondary structure. In* 523ΔVal, *O-P loop Met517 is slightly misplaced from WD, while* 523ΔVal *Thr518 aligns with WT Val519 on α-helix P. Thus, up to Val522 (WT) and Leu521* (523ΔVal) *the secondary structure is conserved, although the aminoacidic identity is different. Finally, from Val522 and WT Val523 on, both primary and secondary match again (only aminoacidic sequence number is shifted due to deletion of Val523).*

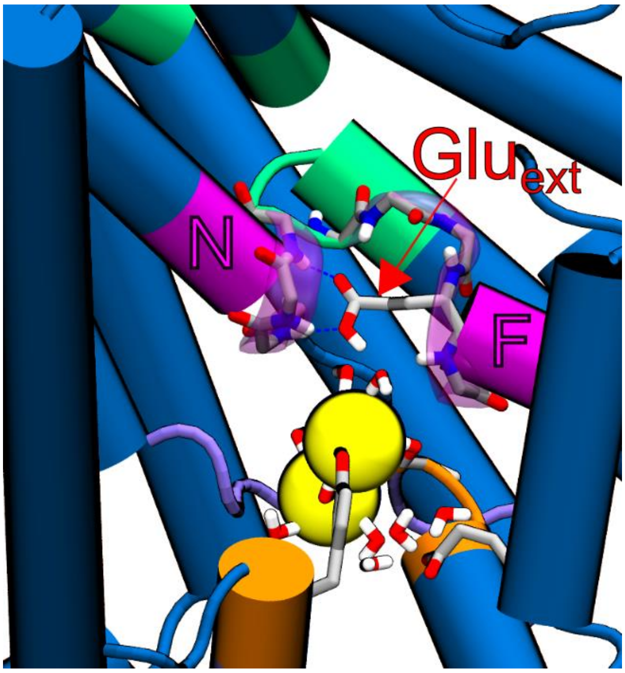


*Figure S3: Interactions of deprotonated Glu$_{ext}$ with NH backbone of helix N, F (including Glu$_{ext}$ NH backbone itself) and residues on E-F loop. This snapshot was extracted from multiple-walker metadynamics simulations, system MP.*

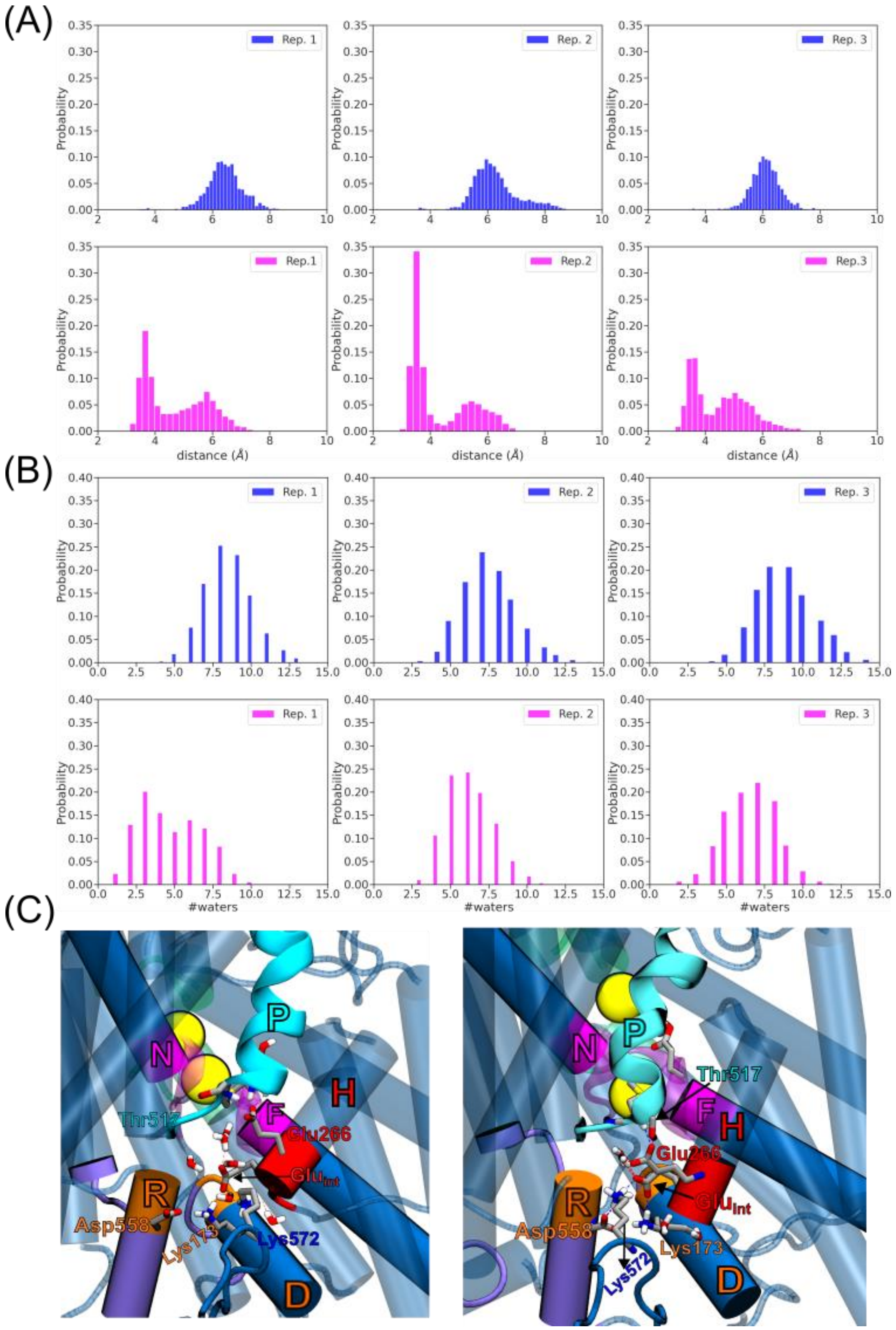


*Figure S4: (A) Distance between Lys172(NH$_3$) and Glu$_{int}$(OE1,OE2) in 3 replicas of WD (top) and MD (bottom) plain Molecular Dynamics simulations. Note that for each replica results from chain A and chain B have been plotted together. (B) Number of water molecules (#waters) around Glu$_{int}$(OE1,OE2) in 3 replicas of WD (top) and MD (bottom) plain Molecular Dynamics simulations. Note that for each replica results from chain A and chain B have been plotted together. (C) Representation of interactions between Thr517 (helix P) and Glu266 (helix H), as well between Lys173 (helix D) or Lys572 (TM-CBS loop) and Glu$_{int}$. Frames have been extracted from M1w and M1m minima.*

### Supplementary Methods

**Definition of the collective variable for MWWTMetaD.** The 2-dimensional collective variable was defined in Plumed[1] as the Z-distance between each of the selected chloride ions and the center of mass of the residues forming the central cavity, $S_{cen}$, chosen as alpha and carbonyl carbon atoms of residues G169, I170, P171, K210, E211, G212, L454, F455, I456, P457, V514, T515, I557 and T558. Due to permutational symmetry, this choice allowed us to augment the obtained data by swapping the two dimensions. In addition, the movement of the ions was restrained in the XY plane to within 6 Å from the respective center of mass of the $S_{cen}$ residues to avoid uncontrolled ion escape close to the top and bottom of the channel. Ions were allowed to sample the Z-distance interval -20 Å < z < 11.5 Å, with semi-harmonic walls with a force constant of 2000 kJ/Å²mol set to all such restraints. In well-tempered metadynamics, Gaussian kernels were deposited every 1000 MD steps with an initial height of 1 kJ/mol, width parameter of 0.3 Å, and a biasfactor of 10. The reweighting factor[2] was calculated on the fly to enable subsequent reconstruction of the free energy surfaces and selection of high-probability frames for the pKa calculation.

**Preparation and equilibration of membrane systems.** CHARMM-GUI Membrane Builder[3] was ued to set up the membrane-embedded system for Gromacs. The standard multi-step protocol including energy minimization and a series of equilibrations with gradually reduced restraints was performed prior to long equilibration, the latter extended to 1 microsecond for each system in order to identify any deletion-induced structural changes not captured by AlphaFold-Multimer.

**Details of the CMIP approach and pKa calculation.** CMIP[4] is a method to calculate classical molecular interaction potential, defined as a sum of the van der Waals and Coulombic potentials around a macromolecule. In the context of pKa calculations, it was be used to compute the relative energetic cost of removing a proton from a protonated residue, and this cost can be translated into the offset from the pKa of an isolated amino acid through the familiar expression:

$$\Delta pK_a = \frac{\Delta E_{cmip}}{2.303RT}$$

The calculation was performed using a suite of in-house Perl scripts that were previously applied for pKa calculation in [5].

**Detection of druggable pockets using MDpocket.** MDpocket[6] is an algorithm based on Fpocket[7] that accounts for the plasticity of the pockets along a molecular dynamic simulation. Unbiased simulations were pre-aligned on the transmembrane region of the monomer A. The last 500 ns of 3 replicas for each variant were used to obtain the map of normalized densities and frequencies of alpha-spheres by following the standard pocket characterization protocol of MDpocket (https://github.com/Discngine/fpocket). Results were inspected using Visual Molecular Dynamics viewer[8] and cavities were identified by slowly increasing the isovalues from the density map until the pocket located between helices P was clearly formed and differentiated. Therefore, the isovalue was set to 5.7 for the WT and 3.3 for the Δ523 variant (Figure 7DEF).
